# Universal Dual-Target CAR-γδT Cells Targeting B7-H3 and IL13Rα2 for Glioblastoma Therapy

**DOI:** 10.1101/2025.05.14.653035

**Authors:** Yang Xu, Yunhao Ye, Yi Wang, Guangna Liu

## Abstract

Glioblastoma (GBM) represents the most prevalent and aggressive primary malignant neoplasm in the adult central nervous system, exhibiting marked infiltrative growth patterns, inevitable recurrence, and dismal therapeutic outcomes with current treatment modalities. While CAR-T cell immunotherapy has demonstrated remarkable success in hematological malignancies, its clinical translation for GBM has been hampered by several fundamental limitations. A key factor among these is tumor-intrinsic heterogeneity, which drives antigen escape and therapeutic resistance. Furthermore, although autologous CAR-T approaches dominate current clinical investigations, they encounter substantial barriers including manufacturing variability, scalability constraints, and practical limitations for widespread clinical deployment. In contrast, allogeneic “off-the-shelf” CAR-T therapy holds greater potential for the future applications. γδ T cells are a particularly compelling candidate for universal CAR therapy, offering several advantages including innate MHC-unrestricted target recognition obviating the need for HLA matching, polyfunctional cytotoxic mechanisms capable of addressing heterogeneous tumor populations, and intrinsic tropism for solid tumors. However, translational implementation has been constrained by their physiological rarity, ex vivo expansion difficulties, and genetic modification inefficiencies. To address these challenges, we adopted a dual-pronged targeting strategy focusing on B7-H3 and IL13Rα2 - two surface antigens demonstrating preferential overexpression across GBM subtypes while maintaining limited distribution in normal tissues. Using phage display platform and function-based nanobody screening we identified high-affinity binders against both targets. Subsequent optimization of γδ T cell expansion protocols and lentiviral transduction parameters enabled the development of a bispecific, allogeneic CAR-γδ T cell platform. Our in vitro studies revealed that dual-target CAR-γδT cells sustained proliferative capacity under GMP-compatible culture conditions, exhibited potent and specific cytotoxicity against antigen-positive glioma cells, and critically, they showed superior elimination of target-heterogeneous tumors compared to monospecific CAR-T constructs. These results establish a robust preclinical foundation for clinical translation and highlight the therapeutic potential of combinatorial antigen targeting coupled with allogeneic γδ T cell engineering to overcome the persistent challenges in GBM immunotherapy.

## Introduction

Glioblastoma (GBM), the most common primary malignant tumor in human brain tissue, typically originates from glial cells or their precursor cells, accounting for approximately 81% of central nervous system malignancies^1^. High-grade gliomas (WHO grades 3 and 4) exhibit an incidence rate of 3-5 per 100,000. The median survival time is 37.6 months for grade 3 and 14.4 months for grade 4 tumors. As the most prevalent primary malignancy in brain parenchyma, GBM constitutes over 50% of all primary central nervous system malignancies^2-4^. Current clinical management of GBM primarily relies on maximal safe resection followed by adjuvant chemoradiotherapy. However, the highly infiltrative and proliferative nature of GBM frequently leads to tumor recurrence postoperatively, resulting in poor prognosis and limited survival. Existing non-surgical therapeutic approaches for gliomas include monoclonal antibody therapies^5^, oncolytic virus therapy^6^, and tumor-treating fields (TFF)^7^, though none have demonstrated broad clinical efficacy. Various anticancer strategies, including novel immunotherapies, are under investigation in preclinical and clinical studies. Notably, CAR-T cell therapy^8^ has shown promising therapeutic potential in clinical trials.

T cells are classified into two subsets based on their T-cell receptor (TCR) composition: αβ T cells (characterized by α and β chain heterodimers) and γδ T cells (defined by γ and δ chain pairing). In human peripheral blood lymphocytes, αβ T cells constitute the predominant population, whereas γδ T cells generally represent only 1-5% of CD3^+^T lymphocytes. Despite their low circulatory frequency, γδ T cells exhibit broad tissue distribution, particularly enriched in epithelial compartments with highest concentrations observed in intestinal mucosa. As a crucial component of the innate immune defense system, γδ T cells demonstrate unique functional properties including MHC-unrestricted antigen recognition and rapid responsiveness to both self and non-self-antigens. These features enable immediate immunological surveillance against conserved pathogen-associated molecular patterns and stress-induced ligands.

Innovative T-cell-based immunotherapies have demonstrated significant potential in tumor immunotherapy. Chimeric antigen receptor (CAR) T cells, engineered through genetic modification to acquire tumor-specific recognition and cytotoxic capabilities, represent a prominent example of this approach^9^. The CAR structure is an artificially designed antigen-recognition receptor, with third-generation CAR constructs currently being the most clinically utilized configuration. These receptors integrate three core components: a single-chain variable fragment (scFv) derived from monoclonal antibodies, a hinge-transmembrane domain, and intracellular signaling modules containing CD3ζ chains^10^. The scFv domain serves as the critical antigen-binding element, comprising antibody-derived variable heavy (VH) and light (VL) chains connected by flexible peptide linkers. The canonical linker structure consists of three tandem repeats of a glycine-serine motif [(Gly)4Ser]3, designated as (G4S)3, which provides optimal spatial flexibility for antigen recognition^11-14^.

GBM exhibits profound tumor heterogeneity, rendering single-target therapies insufficient to eradicate all malignant cells and consequently limiting therapeutic efficacy. Although chimeric antigen receptor (CAR) T-cell therapy has been clinically investigated in GBM targeting antigens including EGFRvIII, IL13Rα2, and HER2, these trials demonstrated constrained clinical benefits with persistent challenges of tumor recurrence and treatment-related adverse events^15,16^. Heterogeneous antigen target expression and post-treatment antigen loss are one of the important causes of tumor recurrence after CAR-T treatment in GBM. In a phase II clinical trial assessing the immune efficacy of EGFRvIII-targeted peptide vaccines, the vast majority (82%) of patients with tumor recurrence showed loss of EGFRvIII expression^17^. Studies targeting IL13Rα2, HER2, and other antigens have also validated this result^18-24^. Therefore, developing dual-target CAR-T cell therapies combining multiple antigens is an important research direction to improve the efficacy and safety of CAR-T in GBM. Most reported CAR-T cell therapies use single-chain variable fragments (scFv) derived from murine or humanized antibodies. However, traditional CAR designs relying on scFv as antigen-recognition domains have inherent defects that may limit dual-target therapeutic development: (1) The murine components in murine/humanized antibody fragments can cause immune rejection in patients, reducing CAR-T persistence in vivo and affecting efficacy^25^; The large size of scFv may hinder dual-target CAR structural integration. When preparing dual-target CARs using traditional methods, concatenating two scFv antigens results in excessively long transgenes that reduce transduction efficiency and increase the likelihood of protein interaction-induced T cell dysfunction. To address these issues, our CAR structure employs alpaca-derived antibody. In 1993, Professor Hamers Casterman and colleagues discovered naturally occurring unconventional antibodies lacking light chains in dromedary camel serum, termed heavy-chain antibodies (HcAbs)^26^ Unlike traditional IgG antibodies, HcAbs lack light chains and the first constant heavy-chain domain (CH1)^27^. The N-terminal region of the homodimeric heavy chain contains a variable domain called the Variable Heavy-chain domain of Heavy-chain antibodies (VHH), which is structurally and functionally equivalent to conventional antibody antigen-binding domains^28^. These fully functional VHH antibodies exhibit high specificity, diversity, and binding capacity comparable to traditional antibodies. HcAbs (75 kDa) are approximately half the size of conventional antibodies (90-150 kDa)^29^. Compared to traditional antibody-derived Fab or scFv - which require complex eukaryotic production systems with high costs^30^, - VHH demonstrates advantages including low-cost mass production in microorganisms and rapid screening through display libraries^31^, along with suitability for adoptive cell therapy^32^. In summary, VHH is suitable as a dual-target CAR antigen-binding domain due to its small size. Introducing VHH into CAR-T cell therapy allows designing CAR structures with higher efficacy and fewer side effects. The advantages include: (1) High genetic homology with humans reduces immunogenicity and immune response risks; (2) Compact size enables dual-target CAR integration and reduces transduction difficulty; (3) Structural advantages may enable VHH to recognize more epitopes than scFv.

Poor clinical accessibility remains another critical factor limiting autologous CAR-T research progress in GBM and other solid tumors. Malignant tumor patients often miss reinfusion opportunities due to disease progression. Under concurrent immunosuppression and high-intensity chemotherapy, patients exhibit reduced quantity and quality of peripheral blood lymphocytes, leading to low success rates in autologous CAR-T preparation^19,33^. αβ T cells recognize antigens and mediate cytotoxic functions through MHC-restricted mechanisms. Given the heterogeneity of HLA across individuals, αβ T cell transfer requires precise HLA matching between donors and recipients, hindering adoptive cell therapy applications. While universal αβ T cell therapy could circumvent this through endogenous TCR knockout to prevent graft-versus-host disease (GVHD), its development is constrained by complex manufacturing processes and unreliable knockout efficiency^34^. These challenges - high costs, time sensitivity, and inter-patient variability - are common limitations of autologous CAR-T therapies, necessitating universal cell therapy development. Among universal cell therapy strategies^35^, CAR-γδ T cell therapy demonstrates particular potential for solid tumor immunotherapy with significant economic value^36^, γδ T cells serve as ideal universal cellular carriers for GBM treatment due to four key advantages: First, as innate tumor-killer cells, γδ T cells mediate tumor lysis through TCRs, NK-cell receptors, and CD16 molecules, partially overcoming tumor heterogeneity-driven immune escape^37^. Second, their MHC-independent recognition mechanism eliminates graft-versus-host responses and bypasses TCR knockout requirements. Third, γδ T cells exhibit strong tissue infiltration capacity, enabling effective control of metastatic lesions. Fourth, they interact with antigen-presenting cells (APCs) and can function as APCs to activate αβ T cells, orchestrating cascade immune responses against tumors^38,39^. However, CAR-γδ T therapy faces challenges: (1) As antiviral innate immune cells^40,41^, γδ T cells demonstrate lower lentiviral/retroviral transduction efficiency compared to αβ T cells; (2) Their low frequency in PBMCs complicates isolation and activation protocols. Addressing these limitations requires optimization of experimental conditions for clinical translation.

In summary, this study will focus on assembling the aforementioned dual-target CAR onto γδ T cells to address both GBM tumor heterogeneity and the cycle/cost challenges associated with autologous CAR-T cell therapies.

## Methods

### Phage Library Packaging

The TG1 strain (Lucigen) was inoculated onto LB plates, and single clones were picked and cultured to an OD_600_ of approximately 2.0 before cryopreservation. The cryopreserved TG1 cells were resuscitated and cultured to an OD600 of approximately 2.0, infected with phages, and activated for amplification, followed by infection with M13KO7 helper phages at an MOI of 20. The cells were allowed to stand for infection at 37°C for 1 hour. After overnight culture, the supernatant was purified by two-time precipitation with PEG8000, adjusted to a titer of 2×10^13^ PFU/mL, and stored.

### Phage library-based Nanobody Panning

Streptavidin-conjugated magnetic beads (Acro) pre-blocked with 3% bovine serum albumin (BSA, Sango) were incubated with phage display libraries and 200–300 nM biotinylated antigens (B7-H3: Novoprotein CY127; IL13Rα2: Sino Biological 10350-H08H-B) at 4°C for 16–18 hours to facilitate antigen-phage binding. After incubation, unbound phages were removed via gradient washing with 1× phosphate-buffered saline containing 0.05% Tween-20 (PBST), followed by elution of antigen-bound phages using acid elution buffer (0.1 M glycine-HCl, pH 2.2).

For binding validation, phage supernatants were analyzed by enzyme-linked immunosorbent assay (ELISA) and flow cytometry. In ELISA, streptavidin-coated plates were probed with phage supernatants, followed by incubation with horseradish peroxidase (HRP)-conjugated anti-M13 antibodies (Sino Biological 11973-MM05T-H) and 3,3’,5,5’-tetramethylbenzidine (TMB) substrate (Invitrogen). Phage clones with a positive-to-negative (P/N) absorbance ratio >5 at 450 nm were considered specific binders. Flow cytometry was performed using PE-conjugated anti-M13 antibodies (Sino Biological 11973-MM05T-P) to quantify cell-surface binding of phages to antigen-expressing target cells.

Positive phage clones were subsequently used to infect log-phase TG1 Escherichia coli cells (OD600 ≈ 0.5). Bacterial pellets were harvested and lysed using Tiangen plasmid extraction kits to isolate phagemid DNA, which was then subjected to Sanger sequencing for clonal validation and stored at -80°C for downstream applications.

### Plasmid Construction

Plasmid construction was performed using homologous recombination. Primers with 15-20 bp homologous sequences at 5’ ends were synthesized by TSINGKE. PCR amplification utilized KOD HiFi DNA Polymerase. Vector digestion employed BamHI and EcoRI restriction endonucleases (NEB). PCR products or digested fragments were separated by agarose gel electrophoresis, purified using Tiangen Gel Extraction Kit (Tiangen), and quantified by OD260. Ligation reactions employed ClonExpress II One-Step Cloning Kit (Vazyme). Ligation products were transformed into DH5α competent cells (Transgene) via 30-min ice bath incubation, 45-sec 42°C heat shock, and 30-60 min recovery before antibiotic plate spreading. Single colonies were screened by colony PCR, with positive clones sequenced for verification. Plasmids were extracted using Tiangen Midiprep Kit and quantified by microspectrophotometer (MiuLab ND-100).

### Cell Culture

To activate human primary T cells, CD3 antibody (Biolegend) was pre-coated at 4°C overnight. Thawed PBMCs (Bokang Bioengineering) were centrifuged at 1500rpm and seeded into precoated plate at 1×10^6^ cells/mL in RPMI 1640 (Thermo Fisher). After 24 hours-activation at 37°C, cells were infected with CAR-carrying lentiviral particles and maintained at 1-2×10^6^ cells/mL for the next expansion. U87 (Hysigen) and other cells were cultured in completed RPMI 1640 or DMEM medium (Thermo Fisher) Supplemented with 10% fetal bovine serum and 1% penicillin-streptomycin (Thermo Fisher). Adherent cells were passaged using 0.25% trypsin-EDTA (Thermo Fisher), while suspension cells were directly centrifuged. Cryopreservation medium contained 10% DMSO, with cells transferred to -80°C via controlled freezing before liquid nitrogen storage.

### Lentivirus Production and T cell Transduction

Lenti-X293T cells were transfected with target plasmid, pMD2.G, pRSV-Rev and pMDlg at an 4:1:1:2 mass ratio by using PEI as transfection reagent. Medium was replaced after 14-16 hours, then viral supernatant was collected at 48/72 hours and concentrated by PEG8000. Viral titers were determined by RFP^+^% in Jurkat cells at 72 hours post-transduction. For human T cell transduction, 5×10^5^ total T cells or 2×10^5^ γδT cells were mixed with concentrated virus (MOI=3-10), and cultured for 72 hours before infection efficiency assessment.

### Overexpression and Knock out Cell Line Engineering

To obtain antigen-overexpressed cells, plasmids incorporating target antigen encoded genes were constructed, and lentivirus were packaged through above mentioned method. The killing curve was established using puromycin (Yeason) at gradient concentrations. Then cells (3-5×10^5^/well) were seeded in 6-well plates for overnight, infected by lentiviral particles, then treated with puromycin at determined concentration. After three cycles of drug treatment, surviving populations were collected and expanded for subsequent experiments. To obtain antigen-deficient cells, CRISPR-Cas9 system-based Knockout was performed. Cells were nucleofected (Lonza 4D) with preassembled RNP complexes (sgRNA:Cas9=100 pmol:31 pmol), with knockout efficiency assessed 72 hours post-transfection. Magnetic Sorting: Cells were incubated with biotinylated antibodies (Thermo Fisher), separated via streptavidin beads (Acro) and MACS columns (Miltenyi), with flow-through (FT) and eluted (ELU) fractions analyzed by flow cytometry (BD). FACS Sorting: Antibody-stained cells were sorted using BD Aria III, with purity verified 72 hours post sorting. All cell lines were confirmed by flow cytometry detection before experiments.

### CAR-T cell Assays

Flow Cytometry detection: Cells were washed with PBS, resuspended in staining buffer with antibodies, incubated, washed twice, and analyzed using BD cytometers. Cytotoxicity: Adherent target cells (8×10^5^/mL) were plated 12 hours pre-assay, while suspension targets were used directly. CAR-T cells were co-cultured with target cells at specified E:T ratios, incubated at 37°C, and analyzed using luciferase kits (Yeason) via Agilent multimode reader. Proliferation: T cell counts were measured every 48 hours using Countstar since Day6 after infection.

### Data Analysis

All experiments were independently repeated at least twice. Flow data were analyzed using FlowJo v10.8.1, sequence alignments with SnapGene v6.0.2, and other graphs generated via GraphPad Prism verion 6, *p* values were determined using one-way ANOVA or two-way ANOVA with Tukey’s correction for multiple comparisons. * p < 0.05, ** p < 0.01,*** p < 0.001, and **** p < 0.0001; ns, not significant.

## Results

### Target selection for dual-target CAR-T therapy for GBM

To address GBM’s high heterogeneity and immune evasion mechanisms, this study employs a dual-target CAR design where antibodies against two distinct targets are serially connected, thereby expanding CAR-T cell targeting breadth^42^. B7 homolog 3 protein (B7-H3), an immune checkpoint molecule with dual co-stimulatory/co-inhibitory functions^43^, demonstrates moderate-to-high expression across GBM specimens while remaining minimally expressed in normal brain tissue^44,45^. Interleukin-13 receptor subunit alpha-2 (IL13Rα2), the first CAR-T therapeutic target validated in GBM, shows tumor-specific overexpression in GBM cells with limited expression in healthy tissues (excluding testes)^46^. Previous studies have independently demonstrated the therapeutic potential of both B7-H3 and IL13Rα2 as promising targets for GBM^3,5,24^. Our bioinformatics analysis of public datasets revealed distinct GBM tumor-selective expression patterns for both targets (Figure 1A). And we observed minimal correlation between their expression levels (Figure 1B), indicating that they may expressed in different tumor populations. Therefore, simultaneously targeting these two biomarkers can cover a broader range of tumor subpopulations, thereby partially mitigating the challenges caused by antigenic heterogeneity and epitope mutation-driven immune escape.

**Figure 1.**
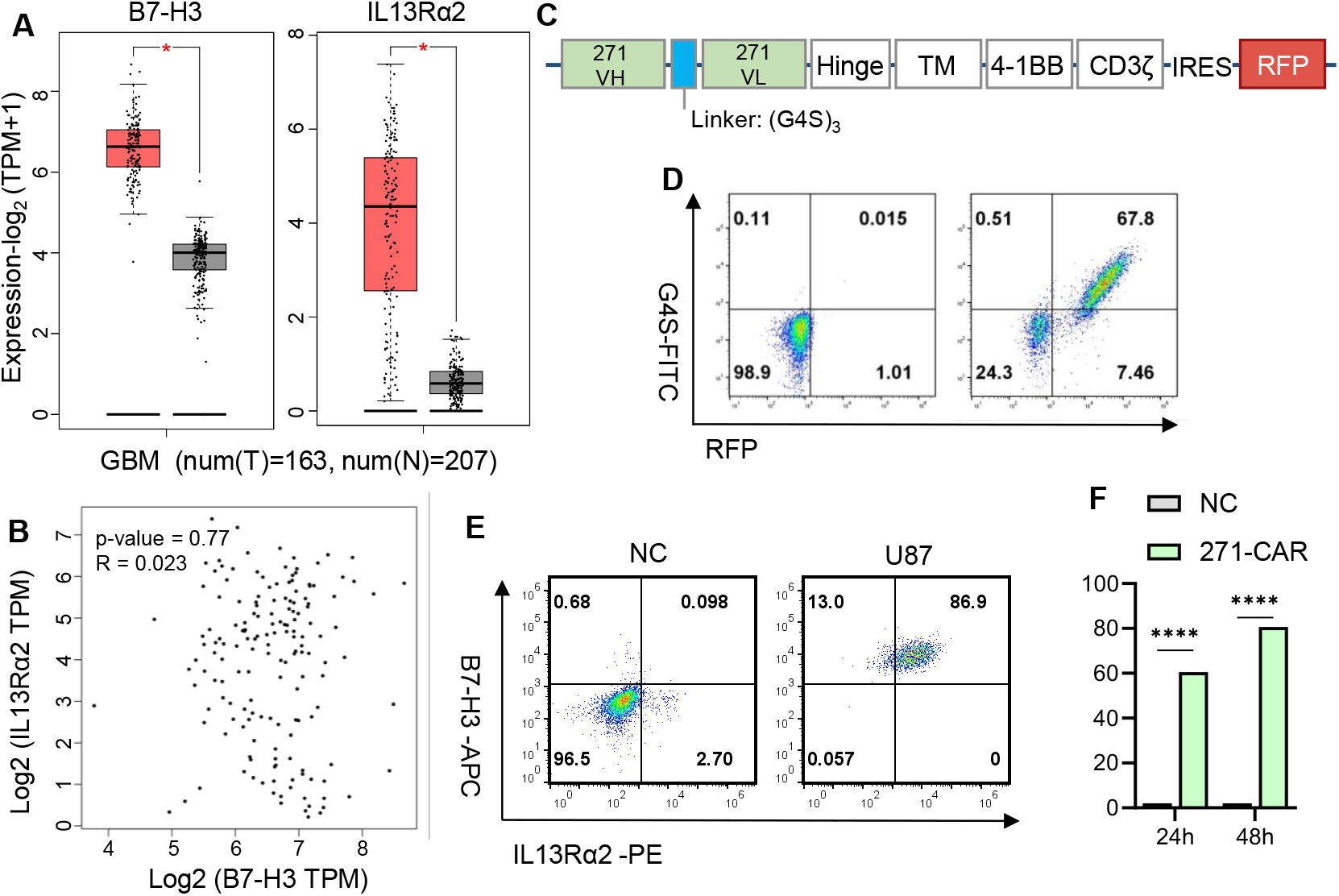
Expression of B7-H3 and IL13Rα2 in GBM Tumor Cells and Killing Efficiency of Single-Target CAR. **A.** Expression of B7-H3 and IL13Rα2 in GBM patients and non-diseased individuals. **B**. The gene correlation analysis of B7-H3 and IL13Rα2 relative expression levels. Original data for A and B are from TCGA database. **C.** Schematic diagrams of MGA271-CAR plasmid structure. **D.** FCM detection of the infection efficiency (RFP+) and the expression of CAR by anti-G4S-FITC antibody on PBMCs. **E.** Detection of the expression of B7-H3 and IL13Rα2 in the U87 cell line by flow cytometer. **F.** Primary human T cells expressing CAR were co-cultured with U87 cells at a ratio of 1:1 for 24 hours or 48 hours, and the death of target cells was detected by luciferase assay then percentage of target cell death was calculated. Data shown as means ± SEM are technical duplicates from one of two independent experiments.

### Establishment of an *in vitro* functional evaluation system for single-target CAR-T therapy

To establish an experimental system for CAR-T cell preparation and cytotoxicity assessment, we utilized functionally validated scFv antibodies incorporated into CAR construct. The B7-H3-targeting MGA271 antibody, with well-characterized functionality^47,48^, was selected for initial validation. VL and VH genes were synthesized and engineered into pCDH-CAR-RFP plasmids (Figure 1C). Lentiviral-packaged construct was used to transduce PBMCs at MOI=3, generating 271 CAR-T cells. Flow cytometry confirmed surface CAR expression (Figure 1D). The GBM cell line U87, expressing both B7-H3 and IL13Rα2 (Figure 1E) was served as target cells. Co-culturing experiments demonstrated 271 CAR-T cells mediated superior cytotoxicity against U87 cells, achieving about 60% and 80% tumor elimination at 24 and 48 hours, respectively (Figure 1F). These results validate the experimental system’s feasibility for functional CAR-T evaluation.

### Identification of specific VHHs via integrated phage display and functional cell screening

Compared to conventional murine or humanized antibody-derived scFvs, nanobody-based VHHs offer distinct advantages for CAR design. To identify B7-H3- and IL13Rα2-specific VHHs, we constructed a naïve phage-based VHH library, and established an integrated screening platform combining phage display library panning (for antigen binding), flow cytometry-based selection (for cell binding), and cytotoxicity assays (for functional validation) (Figure 2A). After two or three rounds of phage panning, successful enrichment was confirmed when phage titers exceeded 10^?^ (Figure 2B). Then 192 monoclonal colonies were randomly selected to validate target antigen binding activity by ELISA. Candidates with higher-antigen binding activity, including 110 B7-H3-targeting VHHs and 144 IL13Rα2-specific clones were identified. Subsequent sequencing eliminated redundant clones with homologous or highly similar complementarity-determining region (CDR3), resulting in 68 unique B7-H3-binders and 94 distinct IL13Rα2-binders. To better mimic physiological antigen-antibody interactions at the cellular level, we implemented FACS-based screening to select VHHs binding to antigen-positive target cells. Eventually, 15 clones specific for B7-H3 (designated B1-B15) and 34 clones specific for IL13Rα2 showed positive target cell binding (designated I1-I34) (Figure 2C). Previous studies found that antibodies selected through conventional phage panning may fail to functionally engage cellular antigens, potentially compromising their utility in CAR-T cell applications^50,51^. Accordingly, we engineered CAR constructs incorporating these VHHs (Figure 2D) for functional validation. Following transduction into Jurkat cells, we observed robust membrane expression in 11 B7-H3-specific and 27 IL13Rα2-specific CAR clones (Figure 2E, 2F). And IL13Rα2-targeting CAR constructs were also transduced into Jurkat-NFAT cells (stably expressing NFAT-inducible luciferase reporter [52]). Eight CAR constructs exhibited robust activation (RLU >200) when co-cultured with Jurkat-IL13Rα2^OE^ cells, while maintaining minimal background activity (RLU <80) against antigen-negative Jurkat controls (Figure 2G). Subsequent functional assessment in primary human T cells revealed that 4/8 IL13Rα2-CAR-T (Figure 2H) and 4/11 B7-H3-CAR-T (Figure 2I) clones achieved >50% specific lysis against U87 cells (co-expressing both antigens) at an E:T ratio of 1:1 for 24 hours. After repeated validation, clone I6 emerged as the superior IL13Rα2-specific VHH candidate, while clone B4 demonstrated optimal characteristics as a B7-H3-targeting VHH, both selected for subsequent studies.

**Figure 2.**
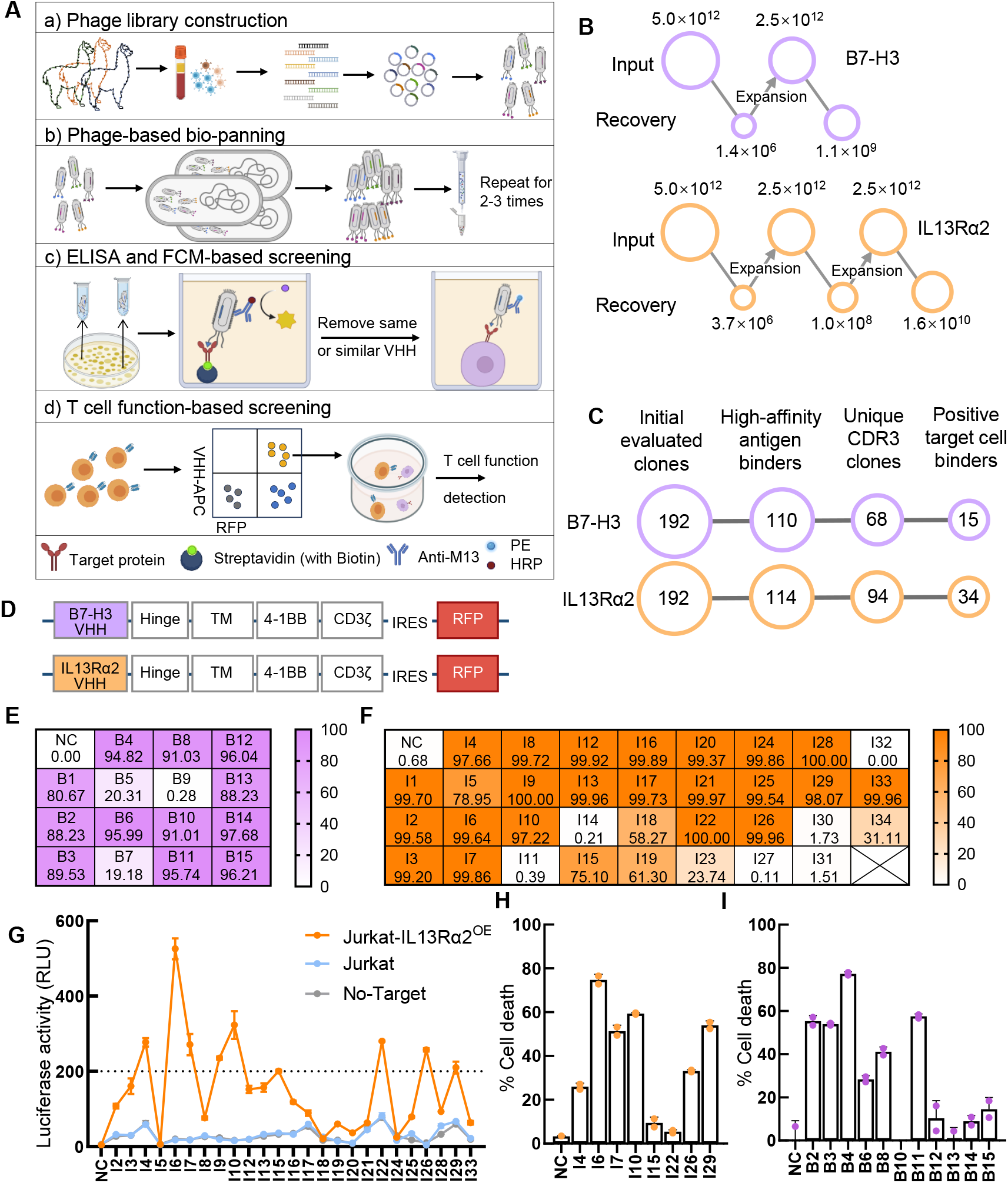
Selection of Nanobodies Based on Phage Display Technology and CAR-T Cell Function. A. Schematic process of phage and cell-based nanobody screening. The nanobody screening process via phage display comprises sequential steps: (a) PBMC isolation from non-immunized alpacas, VHH gene library construction via RT-PCR amplification of extracted mRNA, and phage transformation; (b) Phage enrichment using streptavidin-coated magnetic bead panning, followed by (c) antigen/cell-binding validation via ELISA and flow cytometry (FCM); (d) Engineering validated VHHs into CAR constructs for T-cell transduction, with functional screening to identify high-efficiency clones exhibiting optimal binding and cytotoxic properties. **B.** Changes in the quantity of positive phages in each round of bio-panning. **C. Stepwise screening of** positive phage clones. The numbers after each step-screening were listed as shown. **E, F.** Heatmap of CAR expression profiles on T cells. The CAR expression of 15 CAR-T cells targeting B7-H3 **(E)** and 34 CAR-T cells targeting IL13Rα2 **(F)** was detected by FCM after stained with anti-VHH antibody, the ratio of anti-VHH positive out of RFP-positive populations were analyzed and shown as heatmap. **G.** Jurkat-NFAT expressing IL13Rα2-targeted CAR co-cultured with Jurkat-IL13Rα2^OE^ at E:T=1:1 for 24 hours, and detecting luciferase activity (RLU) via adding substrate. The dashed line shows a cutoff value with RLU=200. **H, I.** The U87 cell death was detected 24 hours after co-culturing with IL13Rα2 **(H)** and B7-H3 **(I)**-targeted CAR-T cells by luciferase assay. Data shown as means ± SEM of two technical duplicates.

### Engineered dual-CAR γδ T cells exhibit superior antigen-selective tumor elimination

For the development of universal bispecific CAR-γδT cells, we first performed systematic optimization of the molecular linkage between two target-specific VHH domains, ultimately obtaining CAR constructs demonstrating both optimal membrane expression and functional activity (Figure 3A). Given the low natural abundance of γδT cells in peripheral blood mononuclear cells (PBMCs, only 0.5%-2%), we established a comprehensive manufacturing platform through rigorous optimization of three critical processes: (1) γδT cell activation, (2) in vitro expansion, and CAR transduction. This optimized platform achieved >80% transduction efficiency for both mono- and bispecific CAR constructs (Figure 3B), along with approximately 2,000-fold expansion by day 16 (Figure 3C). The scalability and robustness of this manufacturing process make it directly translatable for clinical-scale production in future.

**Figure 3.**
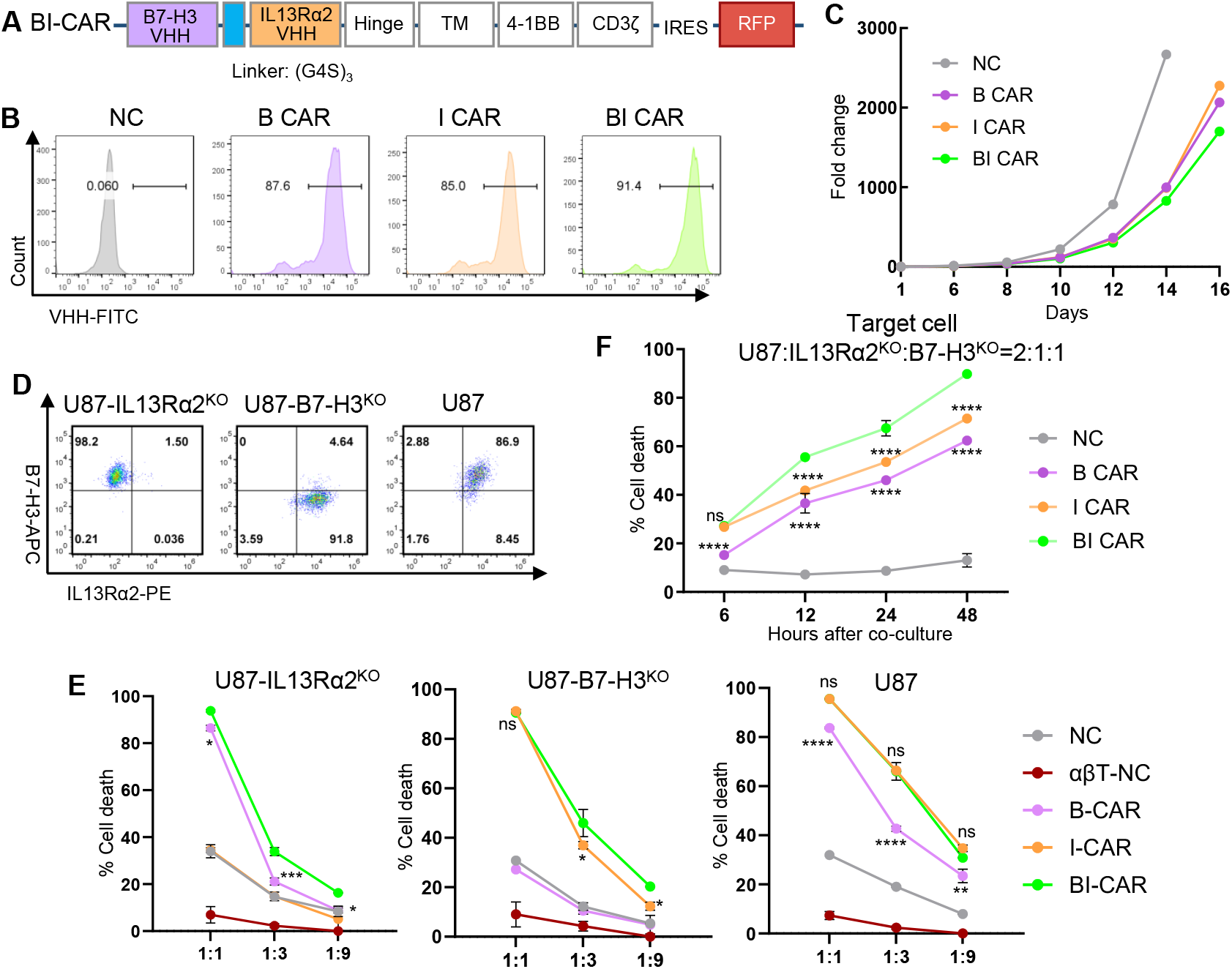
In Vitro Functional Validation of Dual-Target γδ T Cells. **A.** Schematic diagram of the Bispecific BI CAR-RFP plasmid structure. **B.** Expression of B/I/BI CAR in γδ T cells were detected by anti-VHH antibody staining and FCM detection. **C.** Fold change of the quantity of B/I/BI CAR-γδT cells during day1-day16 after activation. **D.** The encoded genes of B7-H3 and IL13Rα2 were knocked out separately in U87 cell line, and single-target cell lines were obtained by flow sorting. **E.** The target cell death co-cultured with single- and dual-target CAR-T cells against U87 cell lines expressing single-antigen and double-antigens were verified in γδ T cells. Untransduced αβ T cells were added as negative control. The cells were mixed at ratios of 1:1/1:3/1:9 and cultured for 24 h, then the target cell death were detected. Data shown as means ± SEM are technical duplicates from one of three independent experiments. **F.** The target cells were mixed at a ratio of U87:U87-IL13Rα2^KO^:U87-B7-H3^KO^ = 2:1:1, then co-cultured with different CAR-γδ T cell at an E:T ratio of 1:3. The target cell death were detected by luciferase assay at 6 h/12 h/24 h/48 h after co-culturing. Three independent experiments were performed. Data shown as means ± SEM of two technical duplicates. * p < 0.05, ** p < 0.01,*** p < 0.001, and **** p < 0.0001, ns: not significant.

To simulate bispecific CAR-T cell activity against antigenically heterogeneous tumors, we created isogenic U87 variants through CRISPR/Cas9-mediated knockout of either B7-H3 (generating U87-IL13Rα2^KO^) or IL13Rα2 (yielding U87-B7-H3^KO^), establishing discrete single-antigen positive cell lines for further studies (Figure 3D). The engineered single- or dual-antigen-expressing U87 cells were employed to evaluate bispecific CAR-γδT cell functionality across effector-to-target (E:T) ratios (1:1, 1:3, 1:9). When co-cultured with single-antigen-positive cells, bispecific CAR (BI CAR) exhibited cytotoxicity comparable to monospecific CARs at high E:T ratios (1:1), but demonstrated superior killing efficiency at lower E:T ratios (1:3, 1:9). Against dual-antigen-positive U87 cells, BI CAR-mediated killing was equivalent to IL13Rα2-specific CAR (I CAR) but significantly exceeded B7-H3-specific CAR (B CAR) (Figure 3E). Notably, untransduced γδT cells exhibited basal tumor-killing activity across all E:T ratios, reflected by their higher basal killing compared to control αβT cells, though this baseline cytotoxicity diminished at lower effector-to-target ratios. This phenomenon likely reflects γδT cells’ intrinsic antitumor activity. These results confirm that the bispecific CAR retains individual VHH antigen-binding capacity and T-cell toxicity. To model tumor heterogeneity, we prepared mixed target cell populations (U87:U87-IL13Rα2^KO^:U87-B7-H3^KO^=2:1:1), creating an experimental model where 25% of cells lacked each target antigen for monospecific CAR evaluation. All CAR-γδ T cell groups exhibited time-dependent tumor elimination, with bispecific CAR (BI CAR) demonstrating superior cytotoxicity compared to monospecific counterparts - achieving 35.21% versus 18.74% (B CAR) and 26.55% (I CAR) killing at 12h, and 89.78% versus 62.41% (B CAR) and 71.42% (I CAR) at 48h (Figure 3F). These results confirm that bispecific CAR-γδT cells maintain potent synergistic activity against heterogeneous tumors while significantly mitigating antigen escape, underscoring their translational promise for clinical applications.

## Discussion

The objective of this study is to develop a universal CAR-T cell therapy with high clinical value for glioblastoma. The antigen recognition domain in CAR molecules, typically comprising scFv (derived from antibody variable regions), serves as the core functional component. Screening antigen-specific antibody sequences remains the most critical step in CAR-T therapy development. We therefore established an integrated technical workflow combining phage display screening and cellular functional screening to efficiently obtain functional VHH targeting B7-H3 and IL13Rα2.

Given the limited genetic cargo capacity of lentiviral packaging systems and the dual-target CAR design requiring two antigen-binding domains, we replaced traditional IgG-derived scFv with VHH from camelid HcAbs. Compared to conventional IgG antigen-binding domains (>400bp), VHH’s shorter gene length (<400bp) significantly reduces lentiviral payload while enhancing transfection efficiency. The compact VHH structure facilitates assembly of dual-target CARs with minimal steric hindrance. Additionally, VHH’s unique architecture enables access to cryptic epitopes inaccessible to conventional antibodies, potentially enhancing CAR-antigen binding affinity and reducing immune escape through disulfide bond formation via exposed cysteine residues. The small VHH size further improves tumor tissue penetration, potentially enhancing CAR-T infiltration in solid tumors like GBM.

We hypothesized that phage-displayed VHH binding might not fully recapitulate CAR-T cellular interactions. Therefore, we incorporated CAR-T-based functional screening to directly evaluate therapeutic candidates. Experimental results validate this approach: phage-validated VHHs showed CAR expression failures (membrane localization defects) and functional deficiencies (equivalent cytotoxicity to untransduced controls), underscoring the necessity of integrated cellular screening in CAR-T development.

Through systematic screening of eight dual-target CAR configurations (D1-D8) with varied linkers and domain orientations, four constructs (D2/D4/D5/D8) demonstrated stable membrane expression and dual-antigen specificity. Functional evaluation revealed D4 (linker2-M288, B4-I6-CAR configuration) exhibited superior T cell activation and cytotoxicity, potentially attributable to M288 linker’s rigid structure enhancing transmembrane signaling. Notably, while all functional constructs showed comparable antigen binding, D4’s activation efficiency improvement suggests linker physical properties (length/flexibility) may modulate CAR conformation and downstream signaling intensity, aligning with reported linker design principles^55^.

The B7-H3/IL13Rα2 dual-target strategy represents an innovative therapeutic approach for glioblastoma (GBM), addressing two critical molecular targets involved in tumor progression. Initiated in 2020, the CAR-T clinical trial by West China Hospital Neurosurgery Department constitutes the world’s first clinical study targeting this combination. GBM’s therapeutic challenges stem from its aggressive nature and immune evasion mechanisms, including antigen loss. While conventional single-target therapies frequently fail due to tumor heterogeneity-driven antigen escape, dual-targeting simultaneously disrupts multiple survival pathways, enhancing efficacy.

Conventional αβ CAR-T cells require MHC-restricted antigen presentation and HLA-matched donors to prevent graft-versus-host disease (GVHD), necessitating TCR knockout with complex manufacturing processes. In contrast, γδ T cells recognize antigens independently of classical HLA molecules, enabling universal CAR-T development without TCR modification. γδ CAR-T approach permits batch production for hundreds of patients and broader tumor applications, overcoming autologous therapy limitations.

Beyond HLA independence, γδ T cells demonstrate superior tumor infiltration. Emerging evidence over two decades highlights their therapeutic potential, including a CD19-CAR γδ T study demonstrating tumor penetration, cytotoxicity, and antigen-presenting cell activation. Addressing γδ T clinical bottlenecks - low expansion efficiency and intrinsic antiviral mechanisms^56^ - we achieved: 1) High-purity γδ T isolation (>95% purity via magnetic/flow sorting); 2) Optimized lentiviral transduction (>90% efficiency with pre-activation); 3) Dual-target synergy showing 2.3-fold improvement over monospecific CARs in clearing antigen-loss variants within heterogeneous tumors.

The BI CAR-γδ T system demonstrates: (1) Broadened targeting scope against B7-H3+/IL13Rα2+/double-positive tumors, reducing antigen escape; (2) “Off-the-shelf” applicability through HLA-independent recognition, eliminating personalized manufacturing delays - particularly critical for rapidly progressing glioma patients.

## Acknowledgements

This work was partially supported by grants from the National Natural Science Foundation of China (82303739), Natural Science Foundation of Hunan Province (2023JJ40198), Hunan provincial department of public education (22A0014, 2024JJ4016).

## Notes

The authors declared that no conflict of interest exists.

### Competing Interest Statement

The authors have declared no competing interest.

